# Patterns of roost site use by Asian hornbills and implications for seed dispersal

**DOI:** 10.1101/2020.09.01.277608

**Authors:** Rohit Naniwadekar, Akanksha Rathore, Ushma Shukla, Aparajita Datta

**Affiliations:** Nature Conservation Foundation, 1311, Amritha, 12^th^ Main, Vijayanagar 1^st^ Stage, Mysuru, Karnataka, India 570017; Centre for Ecological Sciences, Indian Institute of Science, Bengaluru, India 560012; INRAE Val de Loire, Research unit-Forest Ecosystems, Domaine des Barres, Nogent-sur-Vernisson, 45290 France

**Keywords:** communal roosting, Great Hornbill, GPS telemetry, north-east India, seed dispersal, Wreathed Hornbill

## Abstract

Animals spend a significant amount of time roosting. Therefore, understanding roosting patterns and the processes that influence roosting behaviour and roost site choice is essential. Hornbills exhibit interesting roosting patterns with some species roosting communally in large flocks. They are important seed dispersers and patterns of roost site use can have a significant influence on seed dispersal distributions and thereby on plant recruitment. We documented roost site use by four Great Hornbills (*Buceros bicornis*) and one Wreathed Hornbill (*Rhyticeros undulatus*) at a site in north-east India using GPS telemetry. We examined the influence of riverine habitats, nests and foraging range on roost selection. We determined the proportion of seeds that hornbills disperse at roosts and the dispersal distances of seeds dispersed at roosts from the source trees. Through telemetry, we found that roosts of Great Hornbills were generally in forested habitats. Our telemetry data showed that Wreathed Hornbill roosts were close to the river. These results were corroborated by observational data from roost sites where we had regular detections of relatively large flocks of Wreathed Hornbills and occasionally Great Hornbills. The roost sites were not close to the nest sites and were generally within the 95% kernel density diurnal activity ranges. Hornbills dispersed a small proportion of seeds at roost sites. Seeds dispersed at roost sites had almost twice the dispersal distances compared to those dispersed at non-roost sites. This study highlights variation in roost site pattern across individual hornbills and its implications for seed dispersal.

## Introduction

Animals spend a significant part of their life at roosts which are critical habitat for them. Therefore, understanding the patterns of roost site use is essential. The choice of roost sites may be influenced by access to food resources (Johnston-González and Abril 2019) preference for specific habitats (Zoghby et al. 2016), protection from extreme weather (Peters and Otis 2007), predation pressure (Bock et al. 2013, Johnston-González and Abril 2019), parasite avoidance (Rohner et al. 2000), mate selection and anthropogenic disturbance (Peters and Otis 2007). The factors that influence roosting patterns may differ across sympatric species (Peters and Otis 2007). Certain species roost communally, which may accord foraging benefits through information transfer (Ward and Zahavi 1973) apart from enabling energy efficiency (Williams and Du Plessis 2013) and reduced per-capita predation pressure (Eiserer 1984). Most information on roosting comes from raptors (Bock et al. 2013, Watts and Turrin 2017), water birds (Peters and Otis 2007, Jankowiak et al. 2015, Johnston-González and Abril 2019) and birds in temperate regions (Zabala et al. 2012, Jamieson et al. 2016) with relatively little information from tropical birds (Jirinec et al. 2018).

Hornbills are among the largest avian frugivores in Asian and African tropics. Hornbill species may roost as singles (usually breeding males), in pairs, family groups, smaller flocks or in large communal roosts (>50 to over 2000 birds) (Poonswad et al. 2013). At least 26 of the 62 extant hornbill species are known to roost in small flocks or large communal roosts (Kemp 1995, Datta 2001). Multiple Asian hornbill species roost communally (Kemp 1995). The most spectacular communal roosts are those of the Plain-pouched Hornbills (*Rhyticeros subruficollis*) with around 3000 being reported from Malaysia at a single site (Ho and Supari 2000, Kaur et al. 2011). In general, hornbill species of the *Rhyticeros* genus roost in larger flocks (> 50 birds), with 500-1000 Wreathed Hornbills (*Rhyticeros undulatus*) seen at some roost sites in Thailand (Thailand Hornbill Project 2019). A study in north-east India highlighted that hornbills roost on isolated trees in open riverine grassland areas or on cliff faces with lower tree density adjacent to rivers or perennial streams rather than their diurnal forested habitats (Datta 2001). The number of birds in these riverside roosts changed between breeding and non-breeding seasons with smaller numbers in the breeding season as compared to the non-breeding season (Datta 2001). Apart from observations and counts of hornbills at roost sites, there is limited understanding of how individual hornbills make roost site choices. Hornbills may have a selection of roost sites that may be visited at irregular intervals throughout the year (Kemp 1995). A telemetry study on four groups of Southern Ground Hornbills (*Bucorvus leadbeateri*) revealed that birds preferred riverine habitat for roosting, they spent between 1-4 nights per roost per season, and the number of roosts changed seasonally and across the different groups (Zoghby et al. 2016). Similar information does not exist for most other African hornbills. Most of the information on roosting by some Asian hornbill species is from counts of hornbills at roosts.

The detectability and accessibility of roost sites often determine the choice of sites for roost monitoring. Often these roost sites are along the river or in open areas that are more accessible and easily detected, precluding knowledge on roosting at sites that may be in more forested areas. Monitoring roost sites only in the open areas prevents determining whether hornbills prefer to roost in particular habitats (e.g. along the rivers) or close to their nests or foraging areas deeper inside the forest. Telemetry data can provide accurate information on roost site use by individual birds over time. It can be determined if roost site use is influenced by certain habitat features (that may provide them safety from predators) or their nests or their foraging areas by examining frequency and proximity of roost locations to specific sites. Given that hornbills have unique breeding biology with the female incarcerated inside the nest cavity for several months, and the male singly providing food to the female and the chick/s, nest location can also be expected to influence roosting patterns of birds in the breeding season. They may prefer roosting near their nests. On the other hand, given that the hornbill diet is mainly fruits, which are often patchily distributed, hornbills might prefer to roost in or near their foraging areas. Information such as this is currently lacking. GPS telemetry allows us to investigate these questions in greater detail.

Patterns of roost site use may have a significant implication for the critical ecological role that the hornbills play in tropical forests. Forest dwelling hornbill species are called ‘farmers of the forest’ as they are primarily frugivorous and play an effective quantitative and qualitative role in seed dispersal (Lenz et al. 2011, Kitamura 2011, Naniwadekar et al. 2019a, b). They are known to remove a significantly larger number of large-seeded fruits as compared to other avian frugivores (Naniwadekar et al. 2019a), and play a key role in long-distance seed dispersal (Holbrook and Smith 2000, Lenz et al. 2011, Naniwadekar et al. 2019b). While hornbills scatter disperse large quantities of seeds in the forest (Naniwadekar 2014), they are also known to clump-disperse seeds at their nests and roost sites (Kinnaird 1998, Datta 2001, Kitamura et al. 2008). At nest sites, while clumped-dispersal of seeds may offer an initial advantage to hornbill-diet species (Kinnaird 1998), density-dependent mortality of seeds and seedlings in the long run results in negating any potential advantages conferred by clump-dispersal of seeds at nest sites (Datta 2001). Similarly, at the roost site, dispersed seed densities can be an order higher than in perch sites (and almost half that at the nest sites) resulting in clumped seed dispersal (Datta 2001, Kitamura et al. 2008). At the roost sites, although seedlings of hornbill food tree species do establish initially, very few survive beyond a year (Datta 2001, Kitamura et al. 2008). Apart from density-dependent factors, many roost sites are in open riverine areas with significantly lower tree density and canopy (Datta 2001) and are not favourable for the recruitment of hornbill-diet species. Certain roost sites on cliff faces or steep slopes are also not favourable for seedling recruitment, as the seeds roll off and gather in a pile below. Given that hornbills scatter-disperse seeds in the daytime (Naniwadekar 2014, Naniwadekar et al. 2019b) and clump-disperse seeds at roosts, it will be useful to estimate what proportions of seeds are dispersed at roost sites vis-à-vis non-roost sites. Breeding male hornbills disperse a very small proportion of seeds at nest sites and contribute to scatter-dispersal of seeds unlike the incarcerated females (Naniwadekar et al. 2019b). Similarly, if only a small proportion of overall seeds dispersed by hornbills are clump-dispersed at roost sites, then it may not negatively impact the dispersal of tree species, if some of the seeds are clump-dispersed by a frugivore, which is otherwise scatter-dispersing seeds at large distances from the parent plant, thereby enabling expansion of geographic ranges of plants and maintaining genetic connectivity between different plant populations.

Given this background, the broad objectives of our study were to understand the patterns of roost site use by two species of hornbills (Great Hornbill (*Buceros bicornis*) and Wreathed Hornbill) and its implication for seed dispersal. We first describe the number of roosts used by individual hornbills, the frequency of use of different roosts, and the distance between roost sites on successive nights. We then examine whether the riverine habitats (since hornbills are observed roosting on trees in riparian habitats) and nest influence the roost site use. We also examine the distribution of roost sites used in relation to the diurnal foraging range of the individual hornbills. Lastly, we determine the relative proportion of seeds that are dispersed by hornbills at roost and non-roost sites and the dispersal distances of seeds dispersed at roost and non-roost sites. Seed dispersal distances help assess the role of hornbills as long-range seed dispersers and how movements made for roosting influences this parameter.

## Materials and methods

### Study area

We carried out the study in Pakke Tiger Reserve (area: 861.9 km^2^; 92°36_′_–93°09_′_ E and 26°54–27°16_′_N), which is part of the Eastern Himalaya Biodiversity Hotspot, in Arunachal Pradesh state, in north-east India. We tagged the hornbills over two years in the south-eastern part of the reserve, an area dominated by tropical semi-evergreen forest (Champion and Seth 1968, Naniwadekar et al. 2019b). To the south of Pakke is the Nameri Tiger Reserve in neighbouring Assam state and the Papum and Doimara Reserved Forests are to the east and west of the Pakke Tiger Reserve respectively. The Reserved Forests experience significant biotic pressures. Great (*Buceros bicornis*) (2.2–4 kg), Wreathed (*Rhyticeros undulatus*) (1.4– 3.7 kg), Oriental Pied Hornbill (*Anthracoceros albirostris*) (0.6–0.9 kg) and the Rufous-necked (*Aceros nipalensis*) (2.2–2.5 kg) are found in Pakke. The latter is restricted to the higher elevations. IUCN has classified the Great, Wreathed and the Rufous-necked hornbill as ‘Vulnerable’ and the Oriental Pied hornbill as ‘Least Concern’ (IUCN 2019). We tagged five adult, male Great Hornbills and one adult, male Wreathed Hornbill between October 2014 and May 2016. E-obs tags (Model ‘Bird 1A’; e-obs GmbH; Germany) were used to obtain fine-scale movement information on hornbills. The breeding season of hornbills is between March to August. One non-breeding Great Hornbill was tagged in the breeding season (March 2016), and one Great hornbill was tagged in the non-breeding season (November 2015). We only tagged adult males. Since hornbills are diurnal animals, tags were programmed to take locations at 15-minute intervals throughout the day and turn off at night to save tag battery power. Reliable roosting information was not available for one of the Great Hornbills, whose tag was programmed to shut down at sunset and turn on at sunrise. For all the other birds the tag was programmed to shut down at least 45 min after sunset (∼ 19:00 hr IST) and turn on at least two hours before sunrise (∼ 03:10 hr IST), which allowed us to extract information on hornbill roosting reliably. Based on our field observations, hornbills arrive at roosts latest by 17:00–18:00 hr (IST) in June when the days are longest. Additional details on the methods and the study area can be found in (Naniwadekar et al. 2019a, b). The GPS data for this study can be accessed from Naniwadekar et al. (2019c).

We monitored one roost site for 45 days between April to June (breeding season) in 2015 and two roost sites (including the one monitored in 2015) for a total of 211 roost watch days (190 unique days on 21 days both roosts were observed) across breeding and non-breeding season in 2016. One of the monitored roosts was next to the Pakke river. The other roost was on a hill slope in Darlong village, which is on the banks of Pakke River. While the first roost site was 20 m from the riverbank, the other was 370 m from the riverbank. Both the roost sites were located outside the boundary of Pakke Tiger Reserve in the adjoining Papum RF and were close to human habitation. Two-three observers counted hornbills at roost sites between 16:00 to 18:00 hr (IST). We recorded the species, time of arrival and number of individuals.

### Analysis

To determine roosts of hornbills, we calculated mean displacement distances between consecutive time points (for every 15-min interval) between 03:15–19:00 hr and found that displacement distances were least for 19:00 hr for the five hornbills (< 32 m; range across individuals: 17.9–31.2 m). Therefore, we used the location of hornbills at 19:00 hr as the roost location for the day. Due to GPS tag errors, location data was not available for 19:00 hr (and often for few periods before that) for all days for all birds. Mean displacement distances were higher for time intervals before 18:30 hr. Days for which we had obtained data at 18:30 hr, we had received data at 19:00 hr also. Therefore, we used the data only for the days on which we had location information at 19:00 hr. We used the roost data to calculate the mean displacement distances between roosts on consecutive days to determine how far, on average, were the roosts located on successive days for the different hornbill individuals.

We used hierarchical cluster analysis with complete-linkage method implemented through the ‘stats’ package in R to identify the cluster of points that were within 200 m from each other (R Core Team 2019). Our observations at communal roosts of hornbill indicate that often hornbills roost on multiple trees at a single site and the distance between the trees can be around 100 m from each other. After arriving at the roost sites, they also move between individual trees. While individual roost locations may vary, we considered all roost locations within 200 m of each other as a single ‘roost site’, and the centroid of the locations within 200 m of each other was assigned a unique roost site code. We assigned any roost which was > 200 m from each other as a separate roost site. We then determined the number of unique roosting sites used by individual birds and the mean and the maximum number of nights for which the particular bird used the same roost site.

To determine if the nest of the tagged, breeding birds or presence of perennial rivers/streams influenced the choice of the roost site, we determined the distance of the roost site from its nest (in the case of the two breeding Great and one breeding Wreathed Hornbill) and from the river/perennial stream for all the five individuals. While Naniwadekar et al. (2019b) have reported the hornbill home ranges, we wanted to determine if the roost locations were within or outside the diurnal activity ranges of the individual birds. We determined the diurnal activity ranges of individual birds by using the kernel density estimation method using the library “adehabitatHR” in R (Calenge 2006) to generate the utilization distributions of the individual birds. We used the default “href” function as the smoothing parameter (Worton 1995, Watts and Turrin 2017). Utilization distribution is a representation of the relative space use by an individual bird within its entire activity range (Worton 1989). To determine the diurnal activity range of the different hornbill individuals, we used the locations between 05:00 and 17:00 hr for the breeding birds (GH3Br, GH4Br and WH1Br) and the non-breeding individual (GH5NBr) that was tagged in the breeding season (Naniwadekar et al. 2019b). For the non-breeding Great Hornbill tagged in the non-breeding winter season (GH2NBr), we used the locations between 06:00 and 16:00 hr since the sunrise and sunset is later and earlier in winters respectively. Hornbills start arriving at the roost sites up to half an hour before sunset (Datta 2001) and our long-term observations of hornbills at select communal roosts indicate that the birds mostly leave the roost before sunrise. We independently validated these timings with the mean displacement in every 15-min intervals for our tagged birds to confirm that our selected timings coincided with the diurnal activity of the different hornbills. We plotted the roost locations as identified using the hierarchical cluster analysis on these diurnal activity ranges of the hornbills to determine if the roost locations were within the diurnal activity range or outside it.

We followed the method outlined in Naniwadekar et al. (2019b) to estimate the relative proportion of seeds that were deposited at roost and non-roost (other) sites and determine the dispersal distances of seeds that were deposited at roost and non-roost sites. In Naniwadekar et al. (2019b), we have outlined the method that we followed to estimate the relative proportion of seeds that were dispersed at the nest and non-nest sites. A random starting point was selected following the distribution of foraging sightings across the entire day. We excluded roost and nest locations of birds from this starting point selection since they were unlikely to be fruiting trees. We integrated the movement information with the gut passage time data to determine the end location where the hornbill potentially dispersed the seed. If the end location was within 50 m of the roost location for that particular day from which the starting point was selected, then the seed was classified as dispersed at the roost site. In this case, the roost location was the precise, daily roost location and not the ‘roost site’ (which was centroid of all roost locations within 200 m from each other) that was identified using the hierarchical cluster analysis. We used the 50 m buffer around the roost to account for both the GPS error and the typical canopy extent of the large trees which hornbills often use for roosting. We also determined the dispersal distances of the seeds from their random start location. Additional information on the distribution of foraging sightings over time, gut retention times and the analytical framework can be found in Shukla et al. (in press) and Naniwadekar et al. (2019b). We performed all the analysis in R ver. 3.5.3 (R Core Team 2019).

## Results

We had a total of 214 days of roosting data for the five hornbills (Table 1). The number of days of data available for a single individual varied between 19–72 days (Table 1). The roost locations for the different hornbill individuals are shown in Figure S1. Most of the roost sites were inside the Pakke and the adjacent Nameri Tiger Reserves. A few roost sites of GH2NBr were in the undisturbed forested tracts of Papum Reserved Forest outside the Pakke Tiger Reserve, and one roost site (eastern most site) of the Wreathed Hornbill was outside Pakke Tiger Reserve across the Pakke River close to human settlements in the neighbouring state of Assam (Fig. S1). The mean distance between roosts on successive nights for the different Great Hornbills varied between 130–1051 m (Table 2). For the Wreathed Hornbill, the mean distance between roosts on successive nights was 1305 m (Table 2). There was no consistent difference between breeding and non-breeding Great Hornbills (Table 2). However, the maximum distance between roosts on successive nights was greater than 1.18 km for the two non-breeding Great Hornbills but was less than 710 m for the two breeding Great Hornbills (Table 2).

**Table 1.**
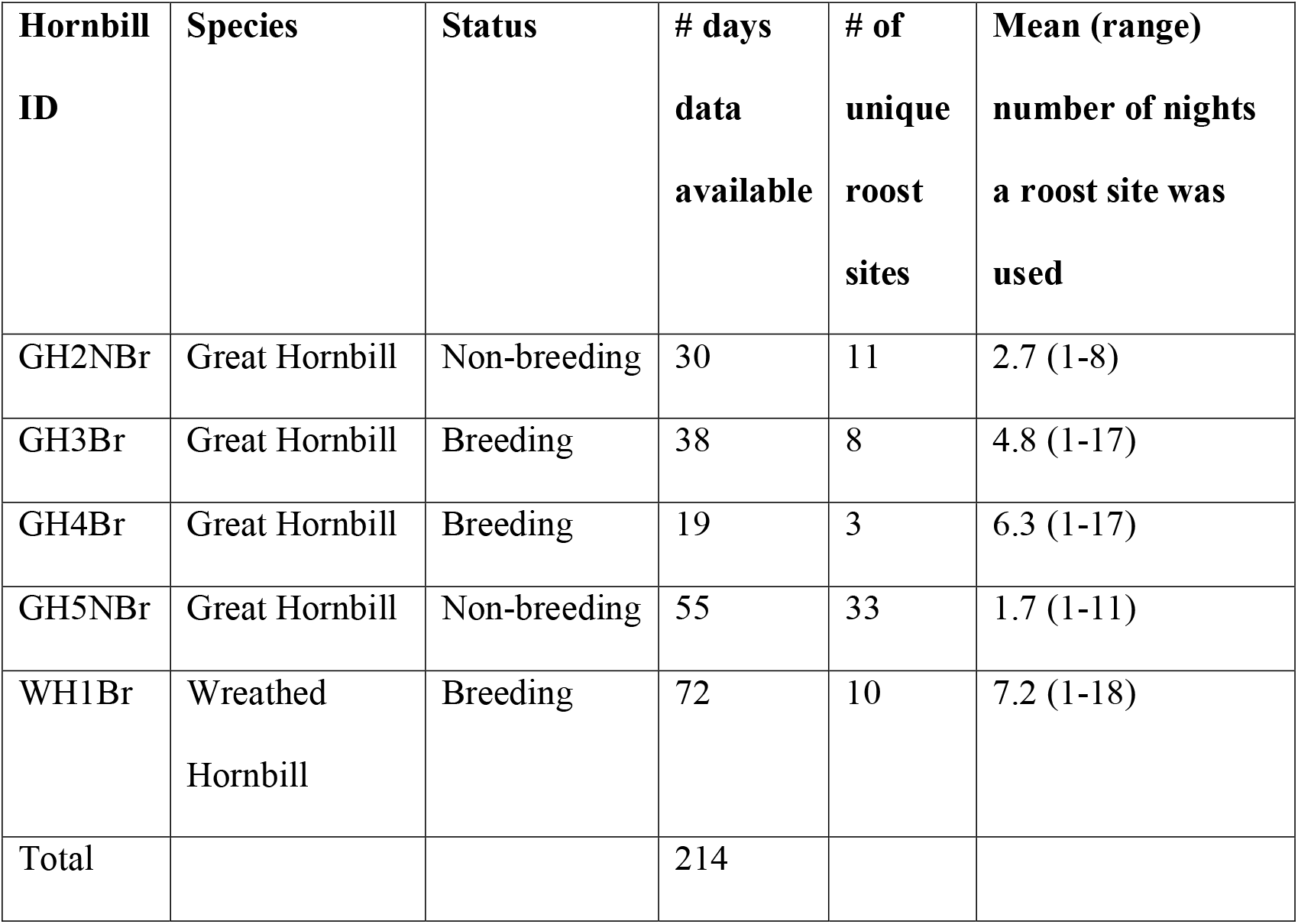
Breeding status, number of days of roosting data available for the five tagged hornbills (one Wreathed and four Great Hornbills), number of unique roost sites (separated by 200 m distance), and mean (range) number of nights a roost site was used by the different individual hornbills.

| **Hornbill ID** | **Species** | **Status** | **# days data available** | **# of unique roost sites** | **Mean (range) number of nights a roost site was used** |
| --- | --- | --- | --- | --- | --- |
| GH2NBr | Great Hornbill | Non-breeding | 30 | 11 | 2.7 (1-8) |
| GH3Br | Great Hornbill | Breeding | 38 | 8 | 4.8 (1-17) |
| GH4Br | Great Hornbill | Breeding | 19 | 3 | 6.3 (1-17) |
| GH5NBr | Great Hornbill | Non-breeding | 55 | 33 | 1.7 (1-11) |
| WH1Br | Wreathed Hornbill | Breeding | 72 | 10 | 7.2 (1-18) |
| Total |  |  | 214 |  |  |

**Table 2.** The average (± SD and range) distance (in metres) between roost sites on consecutive days for the different hornbill individuals and the mean (range) number of consecutive days when the birds used the same roost, and the number of days for which roost data from consecutive days was available is also given. * **–** Number of days for which the roost data was available for successive nights. This number is different from the number of days for which the roost data is available (which is summarized in Table 1) since no roost data was available for some nights during the tracking period.

| **Individual** | **Mean (± SD)**<br>**distance**<br>**between roost**<br>**sites on**<br>**successive**<br>**nights (m)** | **Range**<br>**(minimum –**<br>**maximum)**<br>**distance**<br>**between**<br>**roost sites on**<br>**successive**<br>**nights (m)** | **Mean (range) number**<br>**of successive nights**<br>**when the bird used**<br>**the same roost** | **Number**<br>**of days**<br>**for which**<br>**data was**<br>**available\*** |
| --- | --- | --- | --- | --- |
| GH2NBr | 232.6 (302.0) | 12.6 – 1183.1 | 4.7 (3 – 7) | 16 |
| GH3Br | 327.9 (234.7) | 17.8 – 709.3 | 3.5 (2 – 5) | 30 |
| GH4Br | 130.3 (199.1) | 3.3 – 601.8 | 5.3 (5 – 6) | 18 |
| GH5NBr | 1050.6 (1034.6) | 5.0 – 4318.7 | 2.6 (2 – 5) | 47 |
| WH1Br | 1305.2 (1575.0) | 3.1 – 4698.7 | 3.8 (2 – 6) | 70 |

Hierarchical cluster analysis revealed that the number of roost sites used varied across different individuals for the cluster distance of 200 m (since we classified all points within 200 m from each other as a single roost site; see Methods). For the breeding Great Hornbills (GH3Br and GH4Br), the number of roosts used during the entire tracking period varied between 3–8 respectively (Table 1). There was only 19 days of data available for GH4Br. The maximum number of days for which an individual bird used a roost site over the entire tracking period was 17 days for both GH3Br and GH4Br (Fig. S1A and B). For the non-breeding Great Hornbills, the number of roost sites used during the entire tracking period varied between 11–33 for GH3NBr and GH5NBr (for which data was available for 30–55 days) (Table 1). The maximum number of days for which an individual bird used the same roost site during the entire tracking period was 8 and 11 days for GH2NBr and GH5NBr, respectively (Fig. S1C and D). For the breeding Wreathed Hornbill, we identified ten roost sites during the entire tracking period from 72 days of available data (Table 1). Wreathed Hornbill used two of the ten roost sites for up to 18 days each (Fig. S1E). Both these roost sites were close to the river (Fig. S1E). The mean number of successive nights the five birds used the same roost site (indicating repeated use of the same roost) varied between 2.6–5.3 days (Table 2).

Roosts of Great Hornbills were generally away from the river bank, but those of Wreathed Hornbill were close to the river. The mean (± SE) distance of the roost sites from the river was not very close for the 1850.2 (± 326.2) m for GH2NBr, 3054.4 (± 80.3) m for GH3Br, 941.5 (± 219.4) m for GH4Br, 1536.5 (± 176.4) m for GH5NBr and only 157.6 (± 65.2) m for WH1Br (Fig. 1). The hornbills did not roost near the nests. The mean (± SE) distance of the roost sites from the nest site was 423.9 (± 86.8) m for GH3Br, 964.6 (± 231.4) m for GH4Br and 1754.2 (± 473.5) m for WH1Br (Fig. 2). All the roost sites of the breeding Great Hornbills (GH3Br and GH4Br) (except one for GH3Br) were outside the 50% kernel density utilization distribution (Fig. 2). However, eight of the 11 roost locations of GH2NBr and 16 of the 33 roost locations of the GH5NBr were within the 50% kernel density utilization distribution (Fig. 2). For the Wreathed Hornbill, six of the 10 locations were outside the 50% kernel density utilization distribution (Fig. 2). All hornbills appear to exhibit relatively long bout of flying when they leave their roosts in the morning and when they arrive at their roosts in the evening as was evident by examining the mean displacement at every 15-min interval (Fig. 3).

**Figure 1.**
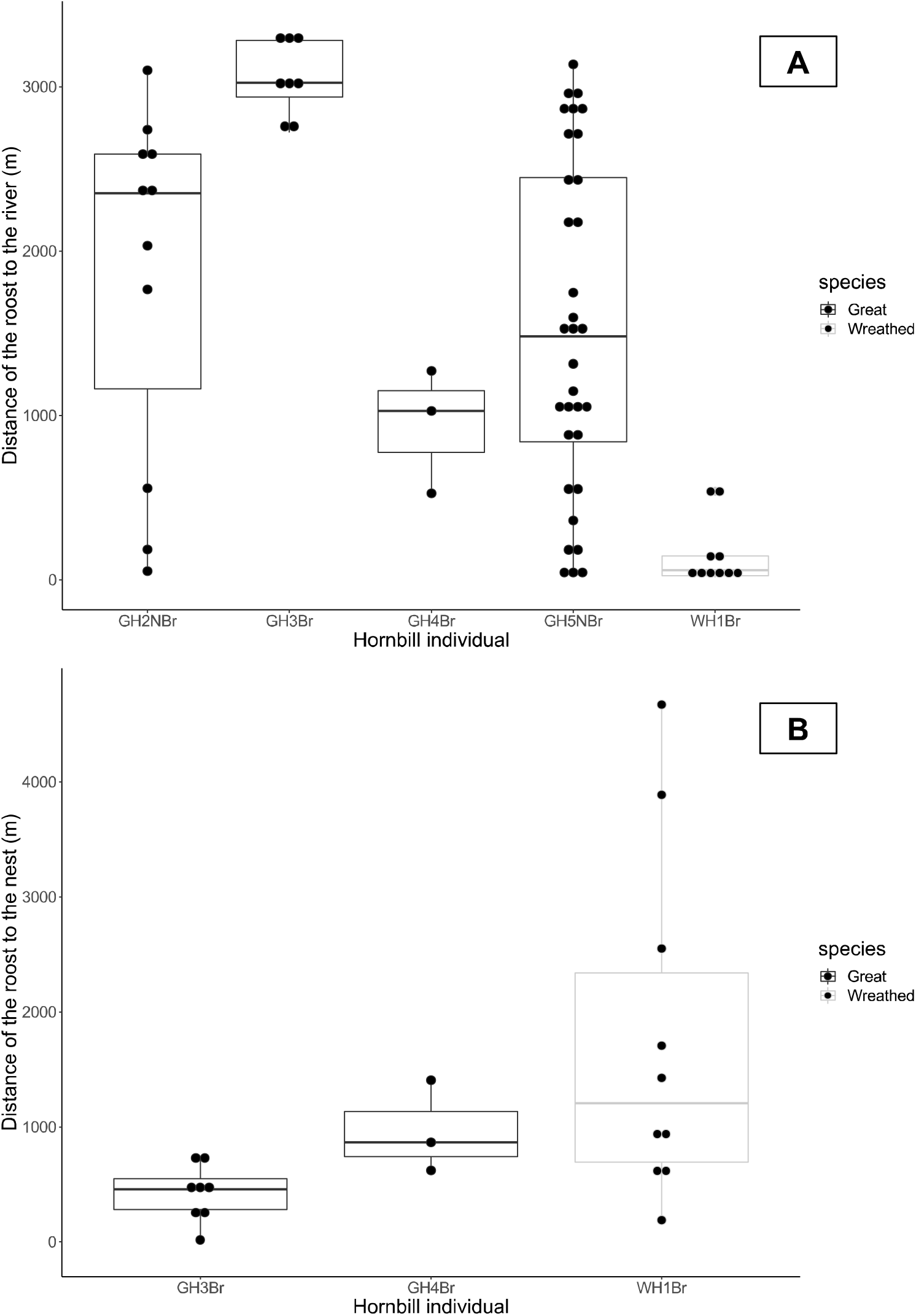
Box and whisker plot depicts that the roosts of the Wreathed Hornbill were close to the river (A) while the roosts of Great Hornbills were not necessarily near the river. The median distances of roosts from the nest sites were above 500 m for the three breeding hornbills. Black-filled points depict individual data points.

**Figure 2.**
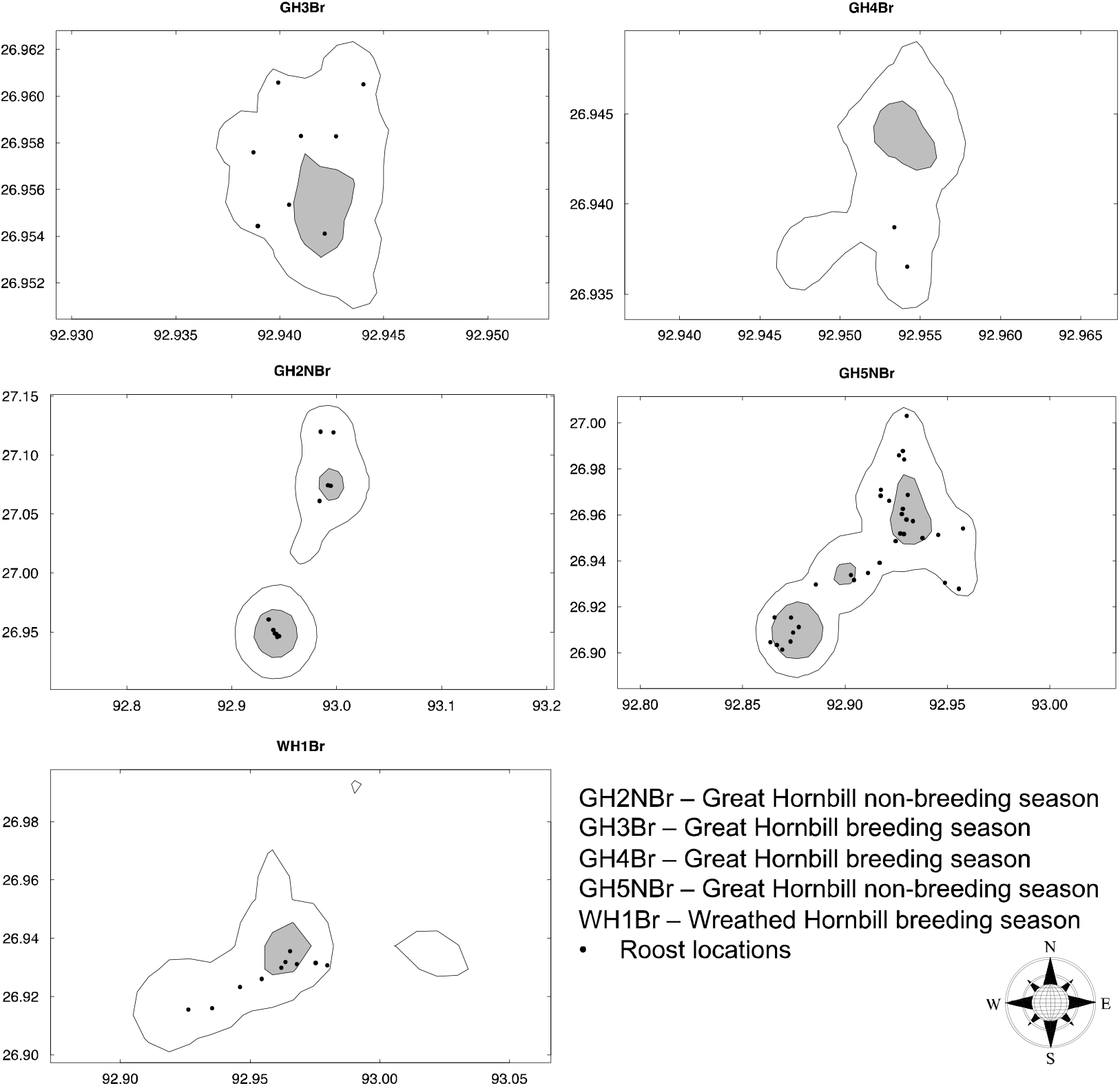
All the roost locations of hornbills (except one roost location for GH4Br) are within the 95% (area enclosed within the black line) kernel density diurnal activity range for the five hornbills but not necessarily within the 50% kernel density activity range (area shown in grey). The locations used for the kernel density diurnal activity range estimation are those between 05:00–17:00 hr for the five hornbills, thereby excluding the roost locations. The black dots are the roost locations of the bird identified using hierarchical cluster analysis. One roost location for GH4Br which was outside the 95% kernel density diurnal activity range is not shown since it was used for only one night and it was far away from its activity range. Coordinates on the map represent the north and east latitudes and longitudes respectively.

**Figure 3.**
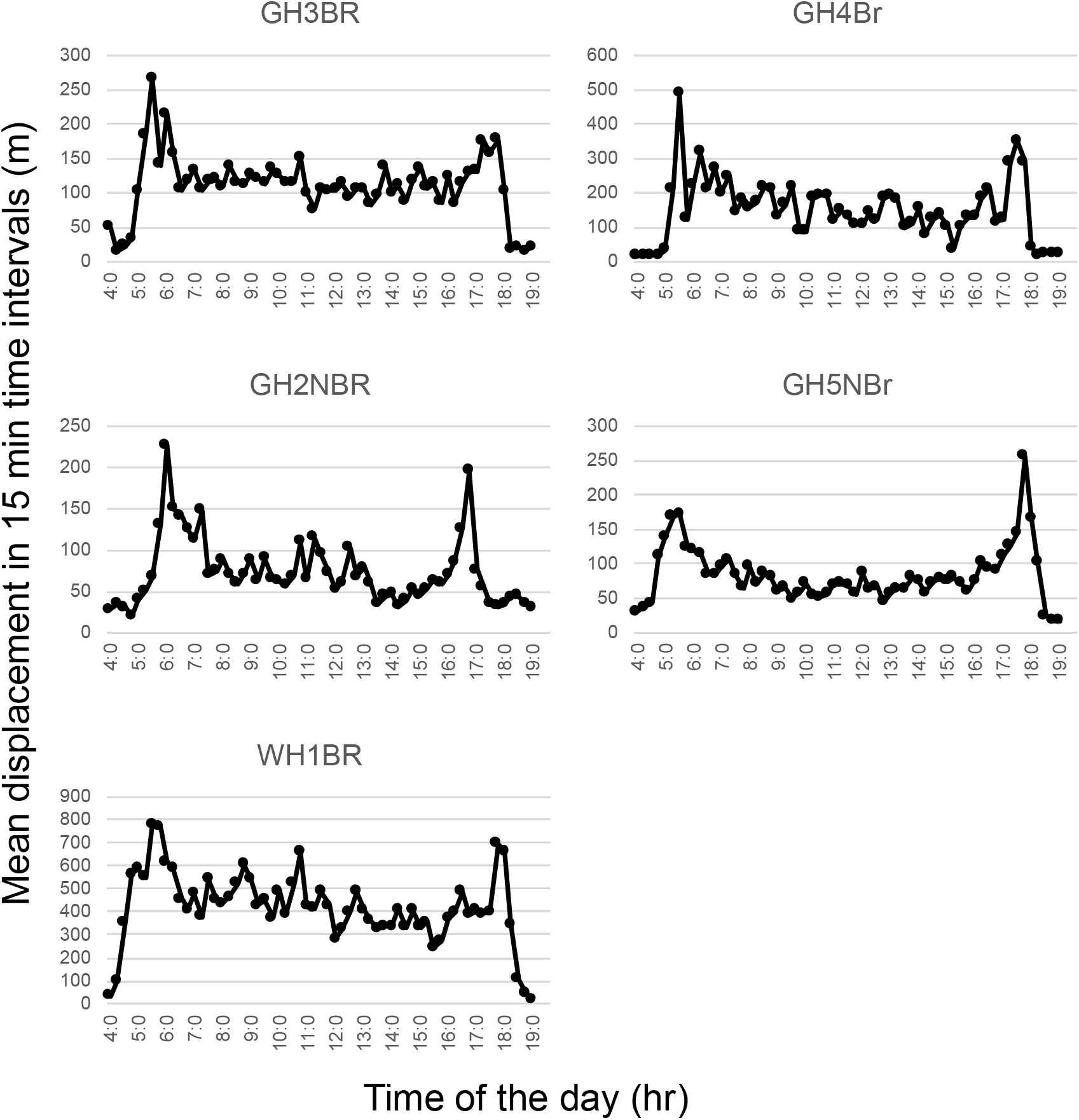
Mean displacement in a 15-min time interval for the five hornbill individuals. GH3Br and GH4Br are breeding Great Hornbills, GH2NBr and GH5NBr are non-breeding Great Hornbills with the former tagged in the winter season (which is the non-breeding season) and the latter in the early summer (which coincides with the breeding season). WH1Br is the breeding Wreathed Hornbill. There is a spike in the displacement just after and before the bird leaves the roost. It indicated that birds travel a long distance after they leave the roost in the morning and before returning to the roost in the evening.

Roost monitoring data indicated that Great Hornbills occasionally (62 out of 256 days of monitoring) used the two riverside roost sites. Great Hornbills were seen on 58 out 157 days of monitoring in the breeding season and four out of 99 days of monitoring in the non-breeding season showing seasonal differences in roost use. The median (range) of Great hornbills when they used the roost site was two (1-5) showing they did not roost in large flocks (Fig. 4). On the other hand, Wreathed Hornbills almost always used the two riverside roost sites (241 out of 256 days of monitoring across the two years). They were seen in 151 of the 157 days of monitoring in the breeding season, and 90 of the 99 days of monitoring in the non-breeding season. Whenever Wreathed Hornbills arrived at the roost sites, they were in relatively larger numbers as compared to the Great Hornbill. The median number of birds was lower in the Darlong roost site (breeding season: median (range) = 10 (1-25); non-breeding season: median (range) = 7.5 (1-17)) as compared to the River Bank roost site (breeding season: median (range) = 25 (1-78); non-breeding season: median (range) = 21 (1-45)) (Fig. 4).

**Figure 4.**
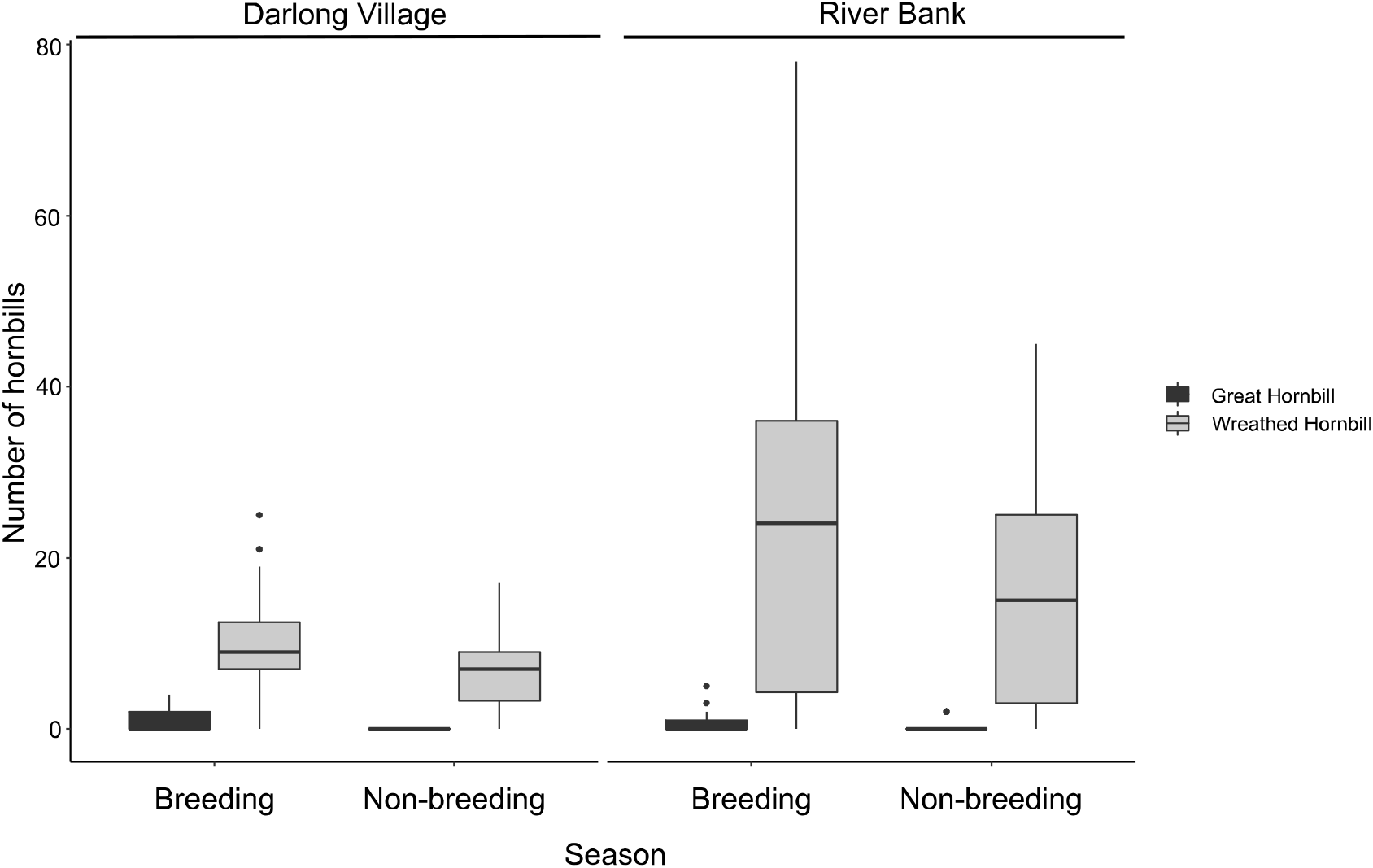
The number of Great and Wreathed Hornbills seen at the two roosts outside Pakke Tiger Reserve in the breeding (March – July) and non-breeding (August – February) season based on data collected by monitoring in 2015 and 2016 based on 161 days of monitoring in Darlong village and 95 days of monitoring at the River Bank site.

The relative percentage of seeds dispersed at the roost sites varied between 7 – 17% (Fig. 5). The breeding hornbills (GH3Br: 7%; GH4Br: 7%) dispersed fewer percentage of seeds under the roost trees as compared to the non-breeding hornbills (GH2NBr: 17%; GH5NBr: 10%) (breeding vs. non-breeding hornbills: χ^*2*^_*1*_=146.6, *P* < 0.001) (Fig. 4). The estimated percentage of seeds dispersed under the roost trees for the breeding Wreathed Hornbill was 9% (Fig. 5). The mean dispersal distances of seeds is higher when they are dispersed at roost sites as compared to non-roost sites (Fig. 5).

**Figure 5.**
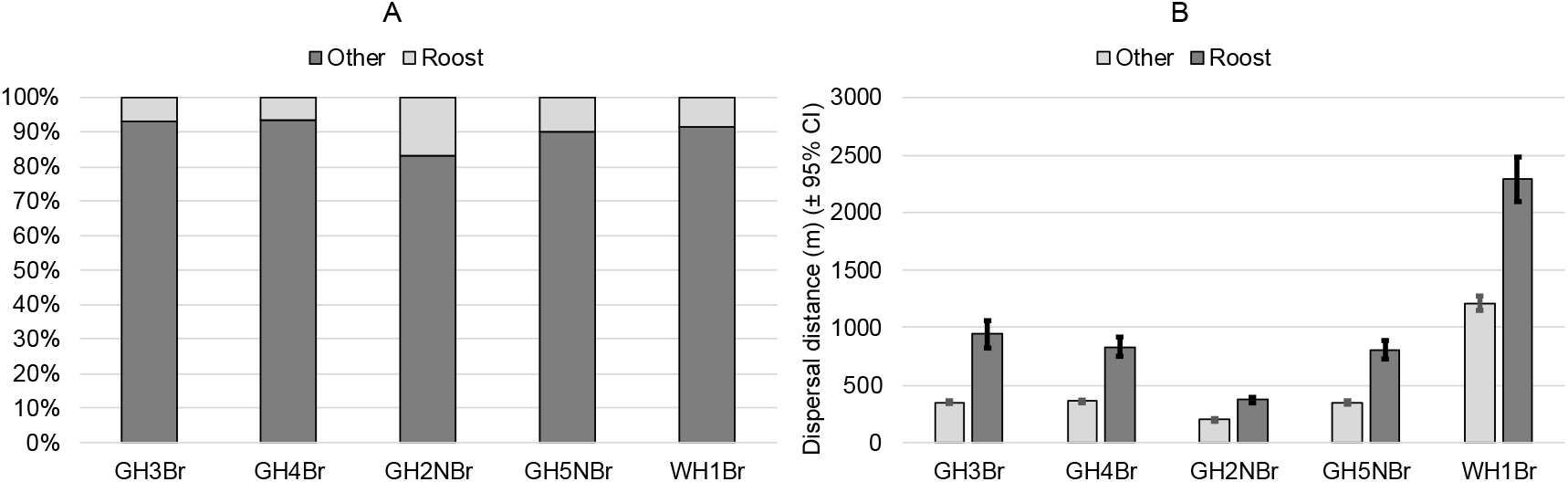
(A) Estimated relative percentages of seeds that hornbill dispersed at the roost and non-roost (other) sites. Most of the seeds are dispersed at non-roost sites. (B) Estimated mean (± 95% CI) seed dispersal distances when the seed was dispersed at the roost and non-roost (other) sites.

## Discussion

This is the first study to examine the individual patterns of roost use by Asian Hornbills and the influence of specific habitat, nest and diurnal foraging range on roost site selection. While Wreathed Hornbills tend to roost near rivers, individual Great Hornbills mostly roost in forested sites away from the river. Both Great and Wreathed Hornbills show some roost site fidelity with individuals using some roosts more often than others. This study highlights that despite exhibiting relatively long commutes to the roost, almost all the roost locations of different hornbill individuals were within the diurnal activity ranges of the hornbills. This study also highlights that hornbills dispersed relatively small proportion of seeds at roost sites. Hornbills dispersed the bulk of the seeds at non-roost sites, which are likely to be more suitable for germination of seeds. Given that individual hornbills use multiple roosts, not all roosts are likely to be used frequently. The infrequently used roost sites might offer favourable opportunities for seeds to establish. Interestingly, the seed dispersal distances at roost sites were more than twice compared to sites where hornbills perch but are not roost sites, facilitating very long-range seed dispersal events during roosting by hornbills. The roost site monitoring data corroborates the finding from the telemetry study demonstrating that Wreathed Hornbills prefer to roost close to the river often in relatively larger numbers as compared to the Great Hornbill.

### Roosting patterns of hornbills

Despite anecdotal reports of several hornbill species that roost communally and in large flocks, there have been relatively very few published studies on the roosting ecology of hornbills (Zoghby et al. 2016). Datta (2001) observed several roosts of hornbills at our study site and reported that Wreathed Hornbills often roosted communally close to rivers or perennial streams and documented seasonal differences in the numbers of hornbills using the roost. Wreathed Hornbills were occasionally accompanied by the Great and by the Oriental Pied Hornbills, mainly in the non-breeding season (Datta 2001). While the Great Hornbills may roost along the river as is evident for the two non-breeding hornbills, our telemetry data for the two breeding Great hornbills demonstrated that they mostly roost away from the rivers/perennial streams in the forests. This is corroborated by the roost monitoring data since we only occasionally saw the Great Hornbills using the roost site which were used regularly by Wreathed Hornbills in relatively large numbers. Whenever they did use the roost site, they did it in small numbers. The two breeding Great Hornbills in our study did not roost near rivers or perennial streams. Great Hornbills range over a very small area in the breeding season (< 2 km^2^) (Naniwadekar et al. 2019b) and the ranges of the two breeding Great Hornbills did not have the river or perennial streams close by which likely explains this pattern of the breeding Great Hornbills not roosting near the river.

Southern Ground Hornbills are known to use multiple roosts and exhibit roost site loyalty (Zoghby et al. 2016). Like the African Hornbills, the two large-bodied Asian Hornbill species use multiple roosts and appear to use at least some of the roosts on multiple occasions. Our limited data indicated that the breeding Great Hornbills seem to have fewer roosts as compared to non-breeding birds. This is expected since the non-breeding areas encompass very large areas (Naniwadekar et al. 2019b) resulting in hornbills roosting in different locations. One of the non-breeding Great hornbills used 33 unique roost sites (sites that were separated by at least 200 m from each other) in 55 days. Despite ranging over large areas, even non-breeding birds appear to repeatedly use some of the roost sites indicating some roost site preference in the non-breeding season.

The role of nests in influencing roosting locations is relatively poorly understood. Given that hornbills exhibit high parental investment in breeding, one can expect nests to influence the roosting of breeding hornbills. However, our data suggests otherwise. In the case of hornbills, GH3Br likely used its nest tree as a roost for a single night only. All the other nights and the other breeding Great Hornbills and Wreathed Hornbill did not roost near their nests. Similar results have been found for juncos (*Junco hyemalis*), where they were found not to roost near their nests (Chandler et al. 1995).

Often the choice of the roosts by birds is positively influenced by the position of the foraging sites (Watts and Turrin 2017, Johnston-González and Abril 2019). The choice of roost sites may also be influenced by factors like thermoregulation (Williams and Du Plessis 2013) and predation pressures (Townsend et al. 2009, Johnston-González and Abril 2019). In some cases, the roost locations may be entirely outside their diurnal activity ranges (Jirinec et al. 2016) or in a completely different habitat (Townsend et al. 2009). If the preferred roost sites are not available within the diurnal activity range, it may entail commutes to and from the roost sites to the diurnal activity range. In the case of the hornbills, the roost locations were not located necessarily in the core of their habitats but were within the 95% utilization distribution values. Hornbills also commuted from and to the roost. The Wreathed Hornbill preferred to roost close to the river as is evident from the distance of the roost sites from the Pakke River. However, given that most roost sites of the Wreathed Hornbills were located close to the river likely explains the movement to and away from the roosts. The riparian habitats, where the Wreathed Hornbills roost, have a distinct assemblage of tree species, many of which are not hornbill food plants (Datta 2001). The mean displacement exhibited by Wreathed Hornbill was more than twice that of the Great Hornbills in the mornings when they left the roost and in the evenings when they returned to the roosts indicating long commutes by the Wreathed Hornbills. However, even during the day, the Wreathed Hornbill ranged over areas larger than the Great Hornbills. Therefore, despite long commutes, the roosts of the Wreathed Hornbills were within the 95% utilization distributions. *Rhyticeros* Hornbills are known to commute relatively long distances to and away from their roosts. Large flocks of the Plain-pouched Hornbills *Rhyticeros subruficollis* are known to fly to and away from the roosts in peninsular Malaysia (Ho and Supari 2000, Kaur et al. 2011). Given that even the Great Hornbill travelled some distances and yet the roosts were within the 95% utilization distributions, suggests that local-scale factors, possibly related to nocturnal predation pressures among others, and evolutionary-scale factors, that potentially influence its roost selection which needs to be examined in future.

Roost site monitoring data corroborates some of the findings of the telemetry data. Wreathed Hornbills regularly roost in relatively large numbers (as compared to the Great Hornbill) on the riverbank site (on *Albizia procera* and *Bombax ceiba* trees) and the village site which is less than 400 m from the riverbank (Fig. S3). While communal roosting would facilitate information exchange, pair formation and accord protection due to dilution effect and greater vigilance, roosting in open, riverine habitats may also accord additional advantage by enabling relatively easier detection of potential arboreal mammalian predators. Additionally, the nocturnal, arboreal predators of hornbills, like clouded leopards and binturongs, are less likely to use open habitats along the river (Grassman et al. 2005, Tan et al. 2017). Great Hornbills, on the other hand, were hardly seen in the riverine or the village site. The abundances of Great and Wreathed Hornbills in the site are similar (Dasgupta and Hilaluddin 2012). Thus, the observed differences in the number of birds also indicates lower preference by the Great Hornbill for riverine sites as is demonstrated by the telemetry data. Datta (2001) reported Great Hornbills communally roosting along the river banks, which we also detected occasionally. It remains to be determined whether increasing disturbance in the riverine areas has negatively affected this pattern.

Hornbills have been reported to roost close to human settlements (Datta 2001). In the late nineties and by 2004, many roost trees in the riverine habitat close to the settlements on the Assam-Arunachal Pradesh border were felled. A single tall *Bombax ceiba* tree that was still standing and being used as a roost site by Wreathed Hornbills, including our tagged Wreathed Hornbill, was felled in 2017. This happened despite local people knowing about the use of this tree as a roost for many years. This highlights the vulnerability of the traditional, communal roost sites, which are often in open areas and close to human habitations, to human perturbations. We observed hornbills starting to roost in the Darlong village outside the Pakke Tiger Reserve towards the end of 2015. It is rare to see hornbills roosting in a village in Arunachal Pradesh, where hornbills are hunted (Naniwadekar et al. 2015). However, in the study area, hunting of hornbills has declined over the years. Roosting of hornbills in Darlong village is an example of how large-bodied hornbills may use human-dominated areas in the absence of direct persecution.

### Seed dispersal at roosts

Hornbills and other frugivorous animals, like primates, have been reported to clump-disperse seeds at the roost sites (Datta 2001, Russo and Augspurger 2004, Kitamura et al. 2008). Often the initial advantage of clump-dispersal of seeds in the form of high seed and seedling densities is negated because of density-dependent mortality factors in the later stages (Datta 2001, Russo and Augspurger 2004, Kitamura et al. 2008). This study demonstrates the variable context of seed dispersal by hornbills at roost sites. This study highlights that hornbills dispersed only a small proportion of seeds at the roost sites. Given that hornbills spend a significant proportion of time foraging away from the roost sites in the daytime, the bulk of the seeds are dispersed away from the nest sites during the daytime (Kitamura et al. 2008). During the daytime, hornbills have been demonstrated to scatter-disperse seeds in large quantities, especially in sites where they occur in large densities (Naniwadekar 2014). Previous studies on clumped-dispersal by hornbills at roost sites have been at known communal roosts of hornbills. However, as this study has indicated that an individual hornbill may not use communal roosts all the time, and they might roost singly and also use certain roost sites less frequently. At such sites, clumped dispersal of seeds is likely to be only because of seeds dispersed by a hornbill over one night. Density-dependent mortality factors for the seeds and seedlings are less likely to occur at these sites, and the probability of seeds to establish would be higher as compared to the communal roosts which hornbills are known to use at least for decades if not more.

Datta (2001) highlighted the unsuitability of the communal roost for the establishment of rainforest tree species given the lower tree density near the riverside roosts and other microsite conditions. However, hornbills may not always roost close to the river as this study has revealed. Telemetry and roost site data on the Wreathed Hornbill also indicates potential inter-species differences in roost site selection. The Wreathed Hornbill was more likely to roost near rivers as compared to the Great Hornbills. Great Hornbill roosts were in the forest sites, often away from the river. At least some of these sites were not communal roosts (based on our long-term field observations on hornbills in the study site), indicating that not all roost sites of Great Hornbills may be unfavourable for seedling establishment.

Interestingly, the seed dispersal distances were almost twice as far compared to those seeds dispersed at non-roost sites during the daytime. This can be explained by the long-flight distances covered by hornbills before arriving at their roost sites. Hornbills are known to mostly carry out long-range seed dispersal (Lenz et al. 2011, Naniwadekar et al. 2019b), however, in the case of roost sites, they appear to carry out extra long-range seed dispersal as compared to the seed dispersal distances during the daytime. Given that not all roost sites might be unfavourable for seed establishment, this long-distance dispersal might be crucial for the maintenance of genetic connectivity between populations of trees and potentially enabling plants to expand their geographic ranges.

Our past knowledge on roosting by hornbills has come from direct observations at communal roosts, however, little was known about the patterns of roosting of individual hornbills. Despite limited sample sizes, this study has generated vital information on the roosting ecology of individual hornbills. This study provides important information on the idiosyncratic roosting patterns of individuals within species and potentially across species. Given that some of the roost sites may be used for decades, the potential reasons for roost site fidelity needs to be identified. This study, along with Naniwadekar et al. (2019b), highlights the context-specificity in seed dispersal patterns and highlights that not all seed dispersal at roosting sites may be of poor quality. In instances, where the bird roosts singly or pairs at roosts that are not regularly used as well as where solitary birds use ephemeral roost sites inside the forest, the quality of seed dispersal provided will not be compromised. The extra-long seed dispersal distance at roost sites has significant implications for plant populations. In future, long-term data on roosting of multiple hornbill individuals is needed to reveal seasonal patterns in roost use.

## Acknowledgments

This work was supported by the Department of Science and Technology (Govt. of India) under Grant No: SB/S0/AS-124/2012 and International Foundation for Science (Sweden) under Grant No: D/5136-2. We are grateful to Mr. Tana Tapi (Field Director, Pakke Tiger Reserve) and his staff for support. We are grateful to the Thailand Hornbill Project team: Pilai Poonswad, Vijak Chimchome, Phitaya Chuailua, Ittiphon Buathong, and Vichai Klinkai for training. We are grateful to Tsongjing Thonger, our field staff Khem Thapa, Tali Nabam, Turuk Brah, Kumar Thapa, Bharat Chiri, Narayan Mogar and Sital Dako, volunteers Malaysri Bhattacharya, Pranjal Mahananda, Gombu Tachang and Sathya Chandra Sagar for assistance in data collection. We thank Jahnavi Joshi, Anand Osuri, Amruta Rane, Sartaj Ghuman, and Soumya Prasad for discussions and help. We thank Goutam Narayan, Nandita Hazarika, Parag Deka and Neelam Dutta for field support. Ethical approval: All guidelines for animal care were followed. Ethics clearance was obtained from Nature Conservation Foundation. We followed established methods of the Thailand Hornbill Project team for the tagging of hornbills. Female and juvenile birds were not tagged. Permission for conducting this research was granted by Ministry of Environment and Forest, (F–No. 1-61/2013 WL), National Tiger Conservation Authority (F.N. 15-5(1)/2014-NTCA) and the Arunachal Pradesh Forest Department (CWL/G/13 (95)/2011-12/Pt./1235-36).

## Data sharing

The data that support the findings of this study are openly available in [Movebank Data Repository] at http://doi.org/10.5441/001/1.14sm8k1d. The complete citation of the data is: Naniwadekar R, Rathore A, Shukla U, Chaplod S, Datta A (2019) Data from: How far do Asian forest hornbills disperse seeds? Movebank Data Repository. doi:10.5441/001/1.14sm8k1d.

## Ethical approval

All applicable institutional and/or national guidelines for the care and use of animals were followed. Ethics clearance was obtained from the Ethics Committee of the Nature Conservation Foundation that gave suggestions that we complied with. We followed established methods of the Thailand Hornbill Project team and consulted senior wildlife veterinarian Dr. Parag Deka, Aranyak to minimize risk to individual birds. Giving primary importance to bird welfare, we did not tag female and juvenile birds. We obtained research and animal capture permits from the Arunachal Pradesh Forest Department, National Tiger Conservation Authority and the Ministry of Environment and Forests, New Delhi and conducted the research under the supervision of the PTR forest officers.

## Author contributions

RN and AD conceived the idea and the study. RN, US and AR conducted the field work and collected the data. RN and AR analysed the data. RN wrote the paper. AD critically revised the manuscript. AR and US gave inputs on the manuscript. This work was part of the grants to AD and RN.

