## Supplementary material for "Patterns of roost site use by Asian hornbills and implications for seed dispersal": Fig. S

SUPPLEMENTAL MATERIAL

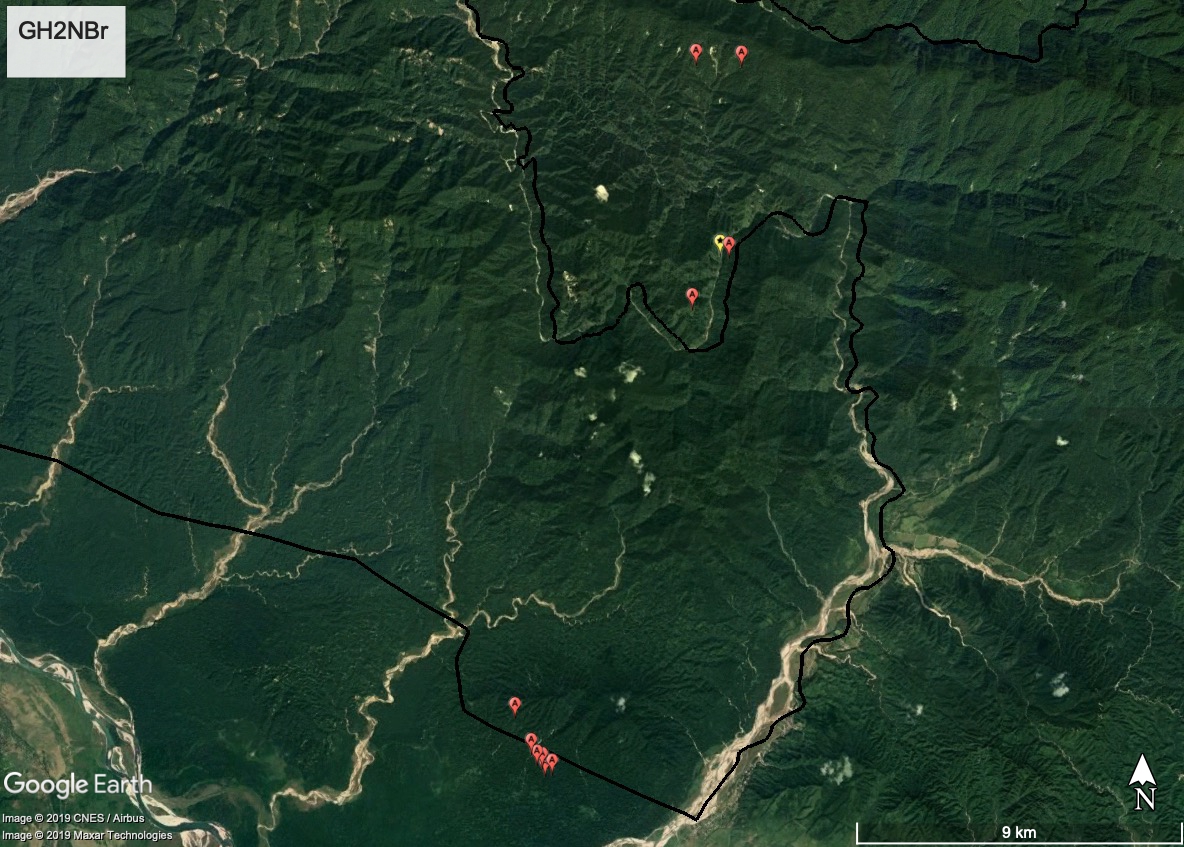

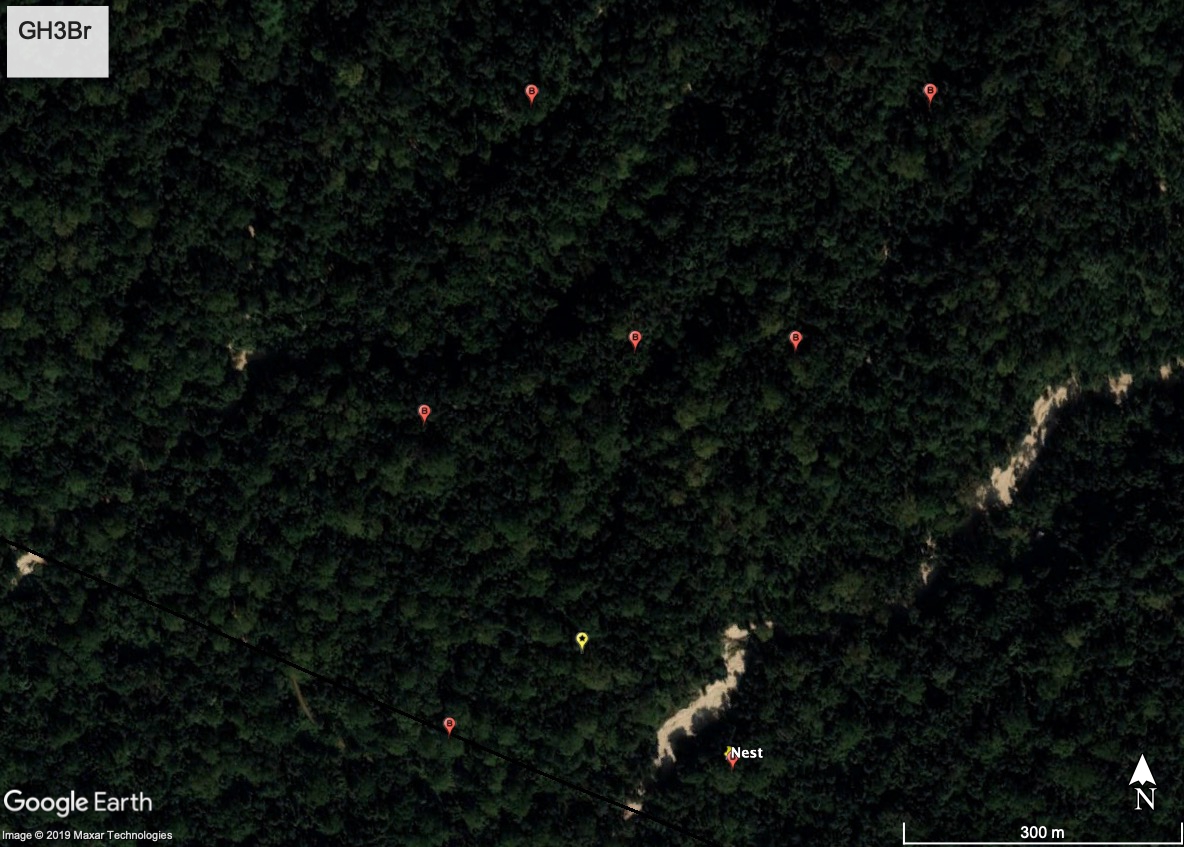

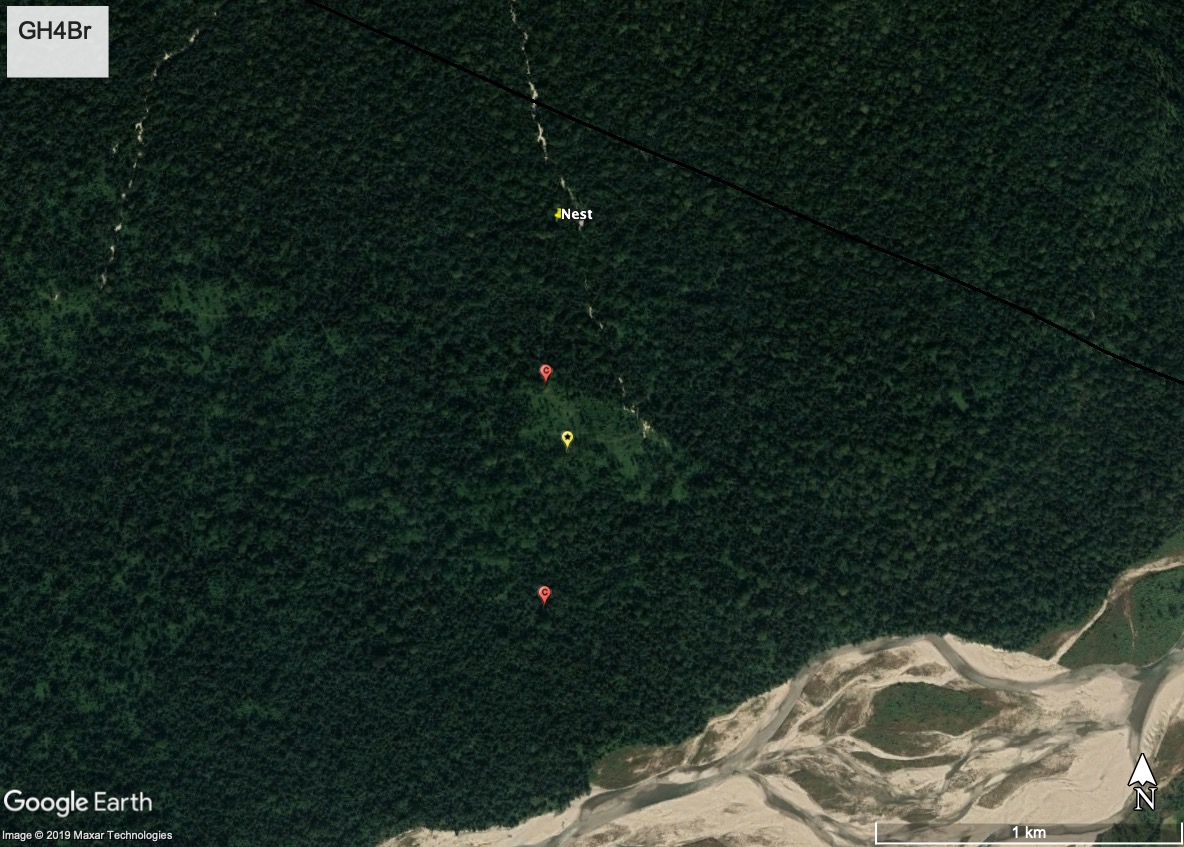

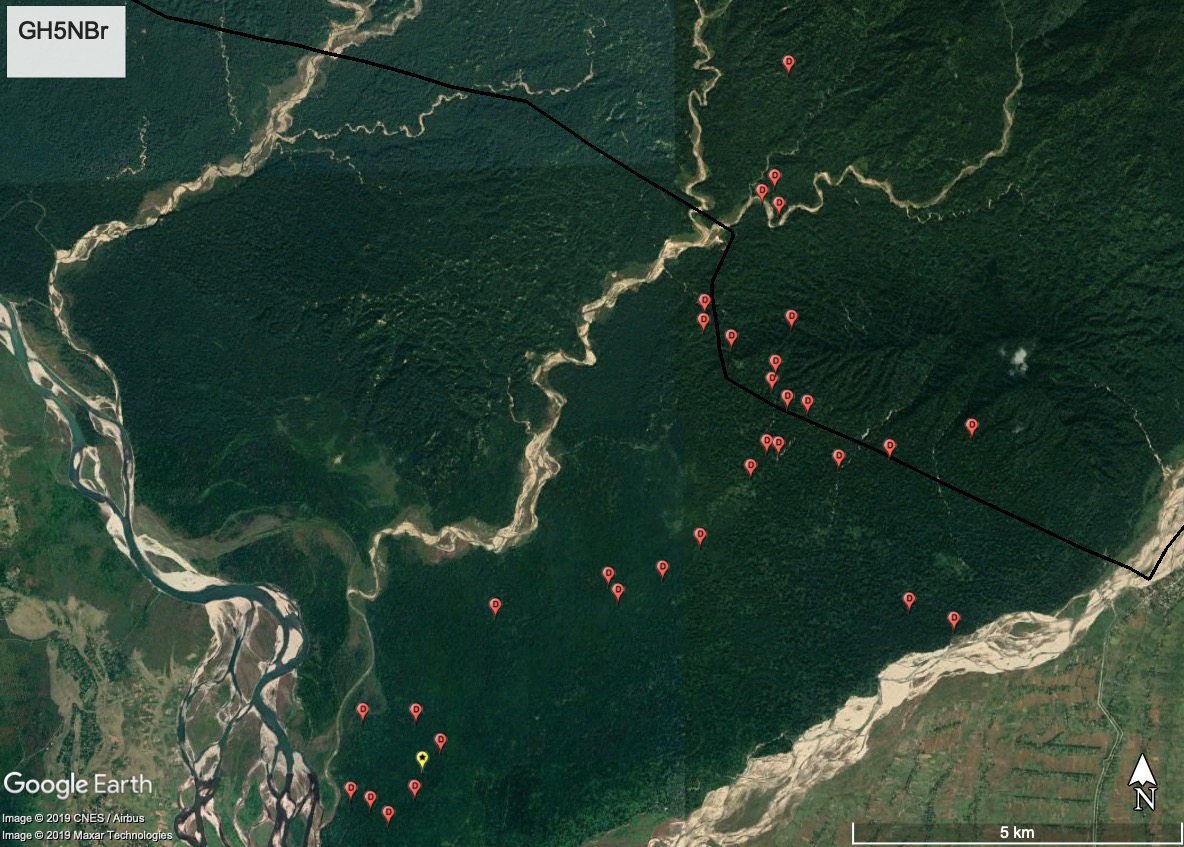

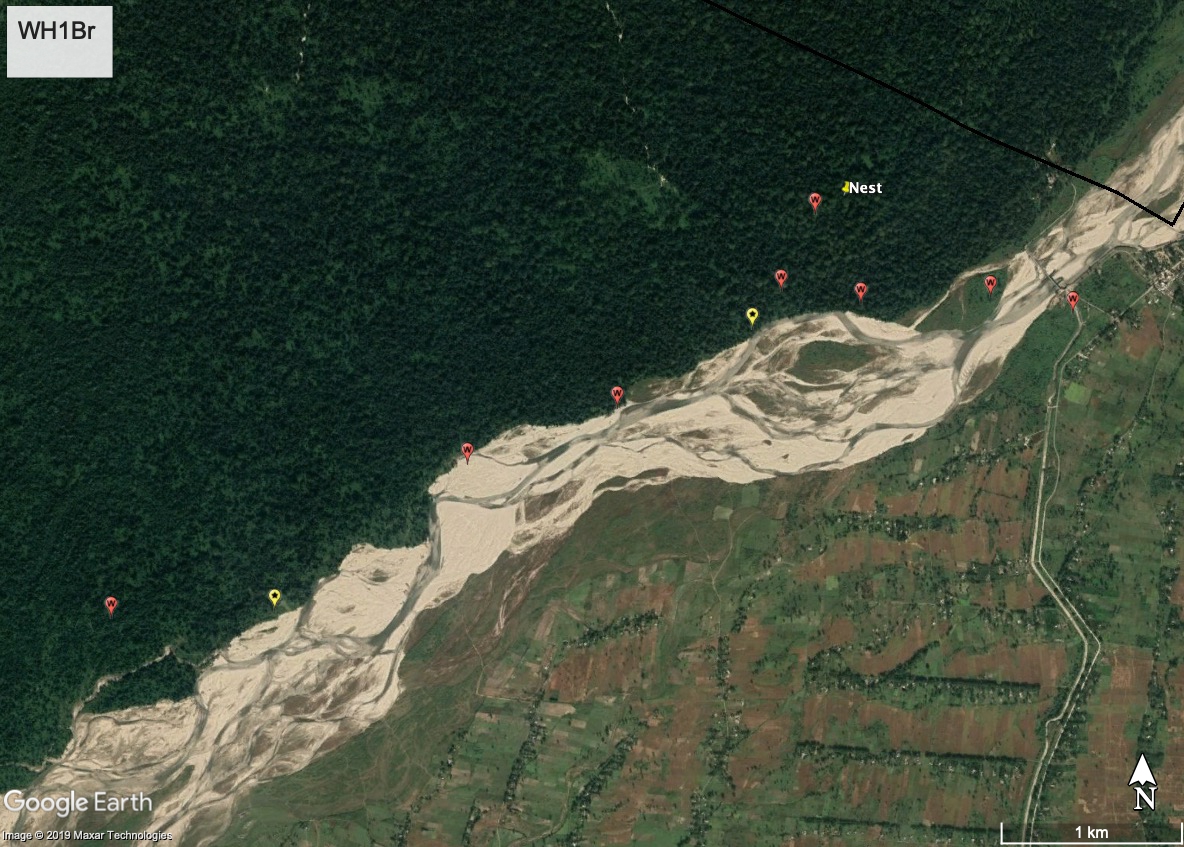

Figure S1. Map showing the roost and nest locations of the five individual hornbills (Great Hornbill non-breeding: GH2NBr and GH5NBr; Great Hornbill breeding: GH3Br and GH4Br; Wreathed Hornbill breeding: WH1Br). The roost locations comprise of all roost sites (across different days) within 200 m from each other and were identified using hierarchical cluster analysis. Nest locations have been marked with a yellow pin. The roost sites most used by the different individual birds have been marked with a star. Area north of the black line is the Pakke Tiger Reserve where the study was carried out and south of the black line is the Nameri Tiger Reserve in the adjoining state of Assam. Since the maps contain information on nest and roost sites of individual birds (some of which are still active), geographic coordinates for the maps have not been provided.

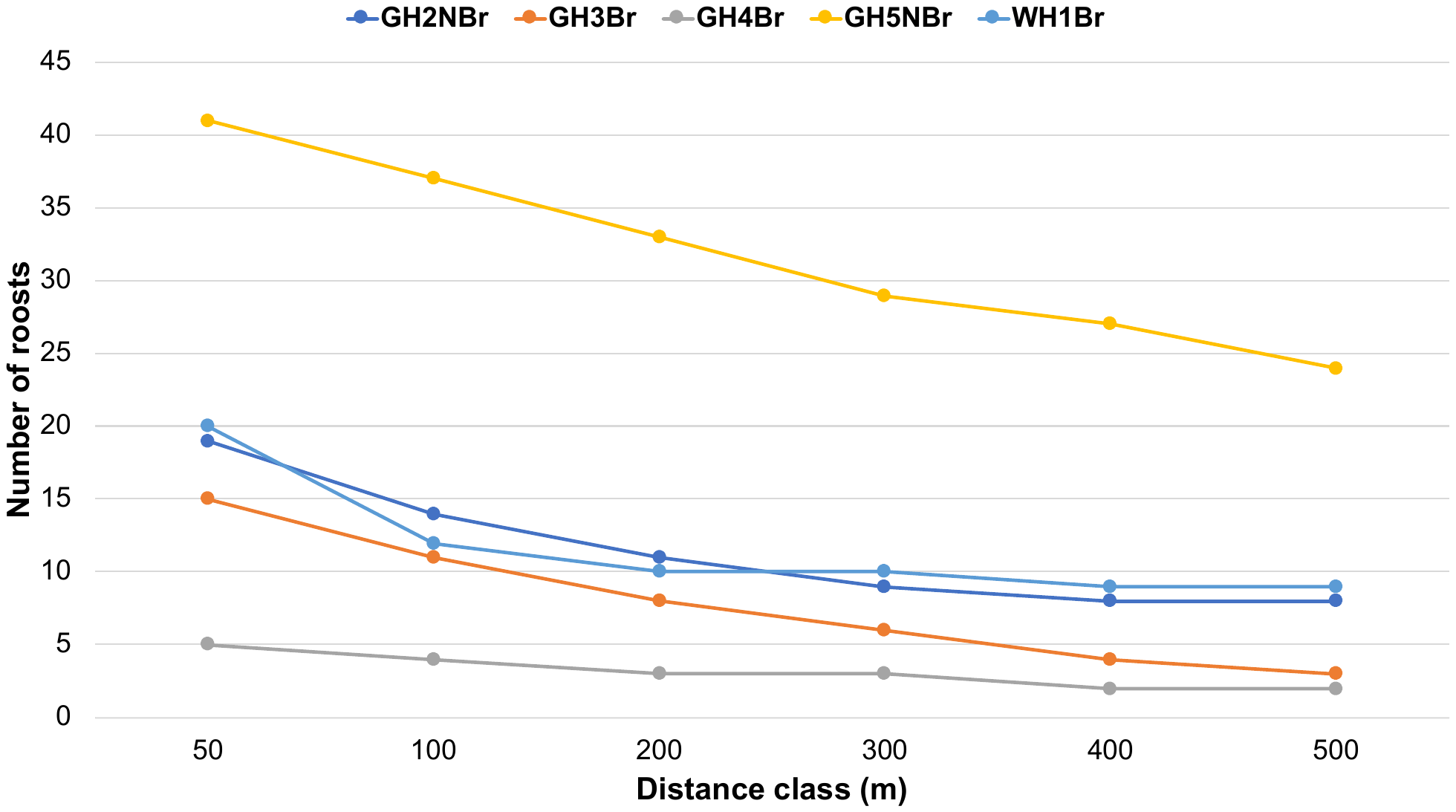

Figure S2. Number of roosts that were identified using hierarchical cluster analysis (complete linkage method) for different distance classes for different hornbill individuals. We selected the distance class of 200 m for identifying distinct roosts for each individual bird.

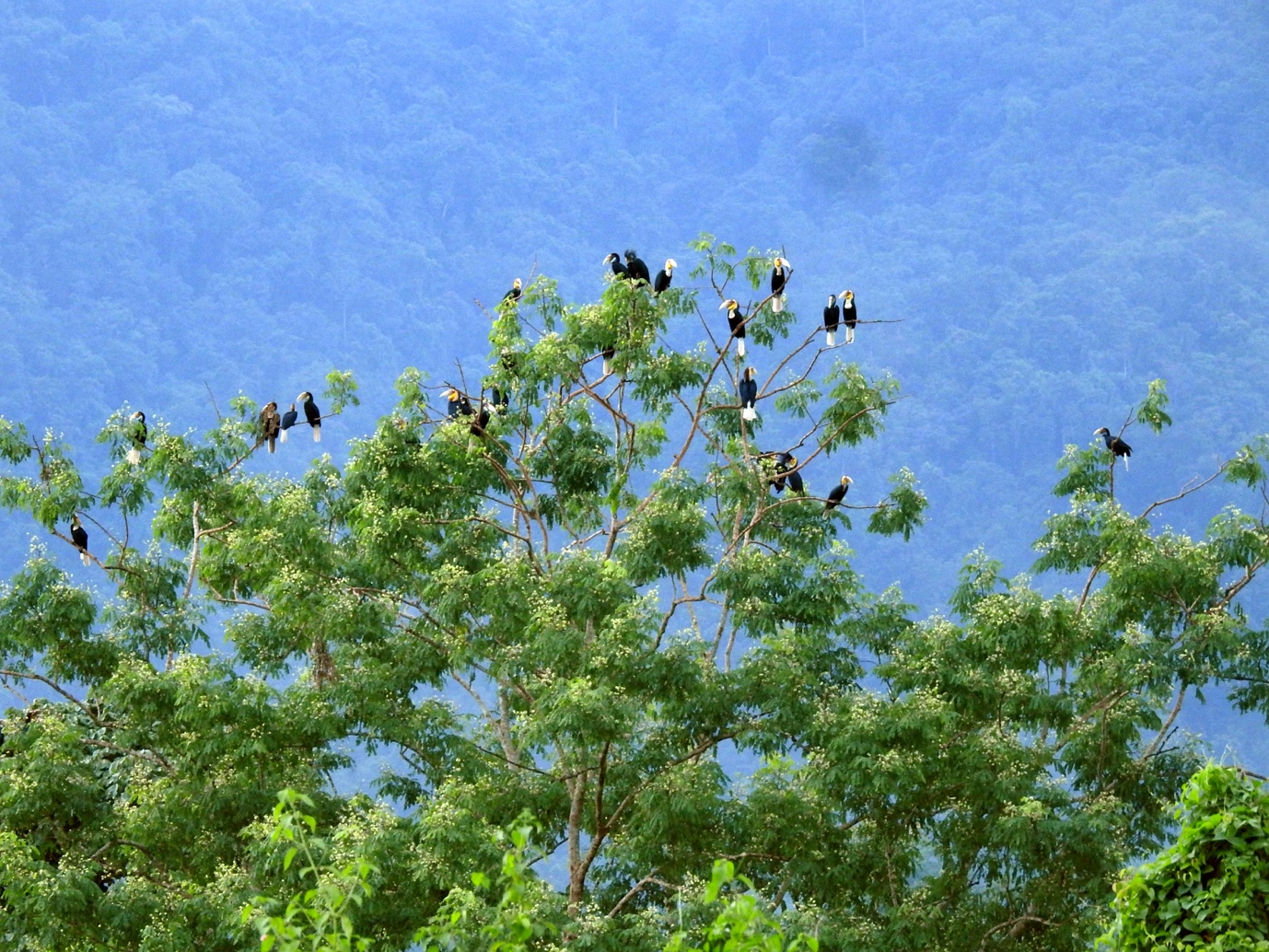

Figure S3. Flock of Wreathed Hornbill *Rhyticeros undulatus* at the roost site on the banks of Pakke River. The roost tree is *Albizia procera*. We monitored this roost site for two years. Photograph by XXXX.
